# Discovering Intron Gain Events in Humans through Large-Scale Evolutionary Comparisons

**DOI:** 10.1101/2024.05.02.592247

**Authors:** Celine Hoh, Steven L Salzberg

**Affiliations:** Department of Computer Science, Johns Hopkins University, Baltimore, MD 21218, USA; Center for Computational Biology, Johns Hopkins University, Baltimore, MD 21211, USA; Department of Biomedical Engineering, Johns Hopkins University, Baltimore, MD 21211, USA; Department of Biostatistics, Johns Hopkins University, Baltimore, MD 21205, USA

## Abstract

The rapid growth in the number of sequenced genomes makes it possible to search for the appearance of entirely new introns in the human lineage. In this study, we compared the genomic sequences for 19,120 human protein-coding genes to a collection of 3493 vertebrate genomes, mapping the patterns of intron alignments onto a phylogenetic tree. This mapping allowed us to trace many intron gain events to precise locations in the tree, corresponding to distinct points in evolutionary history. We discovered 584 intron gain events, all of them relatively recent, in 514 distinct human genes. Among these events, we explored the hypothesis that intronization was the mechanism responsible for intron gain. Intronization events were identified by locating instances where human introns correspond to exonic sequences in homologous vertebrate genes. Although apparently rare, we found three compelling cases of intronization, and for each of those we compared the human protein sequence and structure to homologous genes that lack the introns.

## INTRODUCTION

The discovery that eukaryotic genes are interspersed with non-coding segments, known as introns, was a surprising revelation in molecular genetics when it was first reported in 1977 (Chow et al. 1977) (Berget, Moore, and Sharp 1977). In the decades since, researchers have continued to delve into fundamental questions about introns: their origins, the timeline of their development, and whether their occurrence follows identifiable patterns that could reveal when and how they emerged (Jeffares, Mourier, and Penny 2006; Koonin 2006; Coulombe-Huntington and Majewski 2007; Catania and Lynch 2008; Tarrío, Ayala, and Rodríguez-Trelles 2008; Scott William Roy and Irimia 2009a, 2009b; Ragg 2011; Chorev and Carmel 2012; Scott William Roy and Irimia 2012; Koonin, Csuros, and Rogozin 2013; Wu et al. 2013; Jo and Choi 2015; Lee and Stevens 2016; Catania 2017; Poverennaya and Roytberg 2020). We now know that introns are ubiquitous in plants and animals, with the average number of introns in human protein-coding genes currently estimated at 10.5 (Morales et al. 2022).

The early 2000s saw the first concerted efforts to capture intron gain patterns on a large scale. In 2002, Fedorov and colleagues compared the available data on orthologous animal-plant, animal-fungi, and plant-fungi gene pairs, and found that 39% of fungal introns matched both animal and plant positions, which showed evidence of ancestral introns predating the fungal-plant-animal divergence (Fedorov, Merican, and Gilbert 2002). Another study compared intron structures in 1560 human-mouse orthologs, but did not find evidence of new intron creation within either lineage (Scott W. Roy, Fedorov, and Gilbert 2003). In contrast, a study from the same year that analyzed 684 orthologous gene sets from a diverse range of organisms including animals, plants, fungi, and protists, identified numerous intron insertions in vertebrates and plants (Rogozin et al. 2003).

While those studies were important in advancing our knowledge of the intron evolution, improvements in sequencing technology over the past two decades have greatly increased the number and variety of genomes and the accuracy of their annotations. This progress motivated us to undertake the current study, in which we conducted a large-scale comparison of intron positions between human genes and other vertebrate genes that aimed to discover and determine the timing of intron gain events in an evolutionary context. As the basis for our searches, we utilized the 19,062 proteins in the MANE dataset (Morales et al. 2022), a recently-created high-quality human gene set that contains one “gold standard” splice isoform for each human protein-coding gene. We searched all of these against a comprehensive collection of 17,302,662 proteins from 3493 vertebrate species from the RefSeq database (O’Leary et al. 2016). Our methodology involved identifying introns in every orthologous gene that aligned with a human gene, and then mapping these intron positions onto a phylogenetic tree. This strategy enabled us to pinpoint the appearance of many introns at the base of a specific clade in the tree. Our analysis uncovered 584 recently gained introns in 514 distinct proteins.

Having identified numerous newly gained introns, we then investigated whether we could determine more about their origins. Mechanisms that have been proposed for the *de novo* creation of introns include: intron transposition (Sharp 1985), double-strand break repair (Li et al. 2009), intron transfer (Hankeln et al. 1997), and intronization (Yenerall and Zhou 2012). A more recent study suggested that most new introns are derived from transposable elements, referred to as Introners (Gozashti et al. 2022).

Despite the evidence that a majority of intron gains can be attributed to Introners, we wanted to explore whether intronization might be an alternative method of intron acquisition. Intronization, first proposed in 2008 (Catania and Lynch 2008) (Irimia et al. 2008), is a process whereby part of an exon is spliced out and becomes an intron in subsequent generations (**Figure 1**). Intronization events have been reported within lineages including Cryptococcus (Croll and McDonald 2012; Scott W. Roy 2009), fission yeast (T. Zhu and Niu 2013), Drosophila (Zhan et al. 2014), plants (Z. Zhu, Zhang, and Long 2009), and human retrogenes (Kang et al. 2012). To the best of our knowledge, the only reported instance of intronization across different species groups so far involves humans and chimpanzees (D. S. Kim and Hahn 2012). As of yet, no similar events have been documented within the genomes of vertebrates or extensively compared across their subgroups. We restricted our search for intronization events to cases in which the intron in humans corresponds to an exonic sequence in multiple other vertebrate species. As we describe below, our search identified three novel intronization events in the human genome.

**Figure 1.**
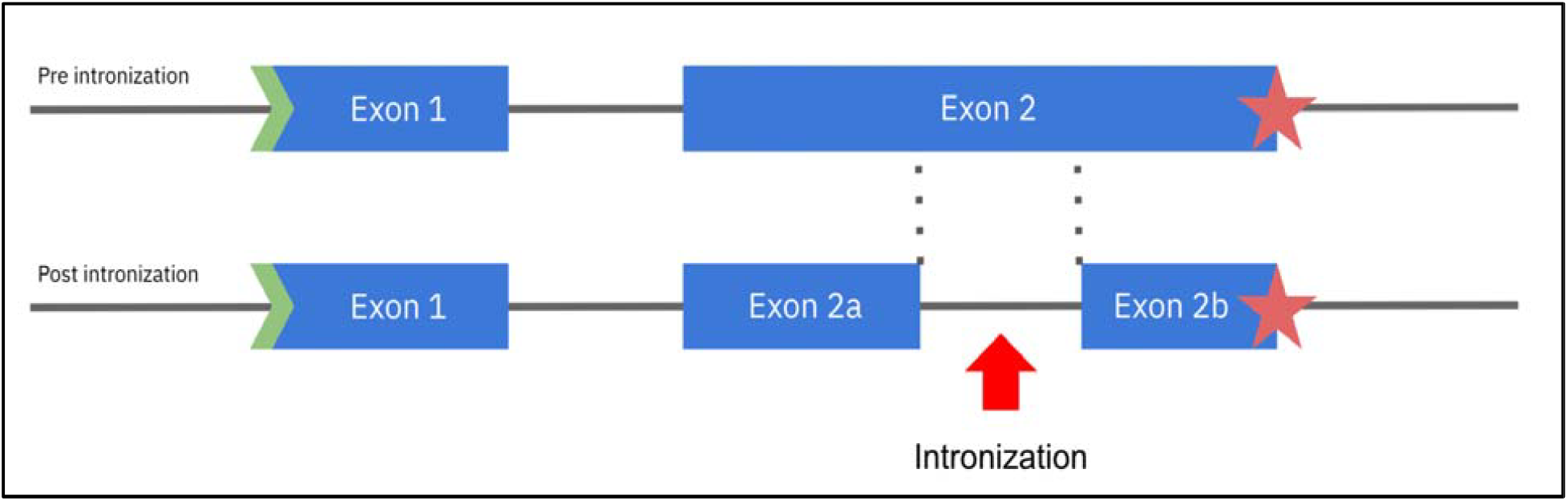
The intronization process. The ancestral form of a gene (top) has two exons, labeled exon 1 and exon 2. After intronization, a segment of exon 2 is transformed into an intron, resulting in the division of exon 2 into two distinct exons, labeled 2a and 2b. The newly gained intron needs to be a multiple of 3 in length in order to avoid causing a frameshift.

## METHODS

### Dataset

We used the MANE (Matched Annotation from NCBI and EMBL-EBI) dataset (v1.0) as our primary source of protein-coding gene annotations (Morales et al. 2022). The MANE database contains a single canonical transcript for each protein-coding locus in the genome, and it was developed with the intent of establishing a universal standard for clinical reporting and comparative genomics. Every transcript in MANE is also contained in the RefSeq (O’Leary et al. 2016), GENCODE (Frankish et al. 2021), and CHESS (Varabyou et al. 2023) human annotations, in which all of the exon and intron boundaries agree precisely among all three catalogs. Release 1.0 of MANE includes transcripts for 19,062 protein-coding genes (plus 58 additional transcripts in the MANE Plus Clinical set), which is over 95% of the total protein-coding gene content in RefSeq, GENCODE, and CHESS. For our database searches, we used the vertebrate subset of the NCBI Protein Reference Sequences (refseq_protein) database, which contained 17,302,662 proteins from 3493 species (as of November 2023). An overview of our analysis process is shown in **Supplementary Figure S1**.

### Identifying intron gain events in human proteins

For each MANE protein, we first performed a BLASTP search (Altschul et al. 1997) against the RefSeq vertebrates database to find all orthologous proteins, keeping the single best hit for each species and requiring a minimum BLAST e-value of 10^-6^. Next, for each human protein and its orthologs, we retrieved gene coordinate tables from NCBI via Entrez Direct (Kans 2024), and used that data to compute the intron positions. We then inserted an “X” character into every amino acid sequence at the position of each intron in that gene. If the intron occurred in the middle of a codon, we inserted “X” just to the left of the corresponding amino acid. We then used MUSCLE (Edgar 2004) to create multiple alignments for each human protein sequence and its orthologs.

In the resulting multiple alignments, we looked for intron conservation focusing on the positions of the human introns. For each intron in each human protein sequence, we assessed how many orthologous proteins had an intron at the same position based on the aligned “X” characters. We then used the NCBI taxonomy to determine where the intron had first appeared. To be considered as a potential recently gained intron, a specific human intron had to be unaligned (e.g., missing) in at least 70% of all orthologous species.

### Analysis of potential recently-gained introns

After obtaining an initial set of intron gain events, we plotted the intron counts against (1) divergence time from humans, (2) intron lengths, and (3) relative intron position (computed as intron position divided by total protein length). We also employed PANTHER to classify the proteins that have gained introns according to their molecular function, biological process, cellular component, and protein class. To identify putative intronization events, we compared the exonic sequences flanking the gained introns in the human proteins with their counterparts in orthologous proteins. If intronization occurred, this comparison should reveal an string of amino acids where the exon (now an intron in humans) was previously located. To detect such patterns, we conducted separate BLASTP searches for the sequences of the two human exons flanking the intron, searching against all vertebrates except for the subgroup containing the intron (**Figure 2**). We retained hits from these searches if they exhibited a gap between the two matching regions in another genome. These hits were considered potential intronization events. Subsequently, we aligned the amino acid sequences using MUSCLE and BLASTP to visualize the evidence of intronization.

**Figure 2.**
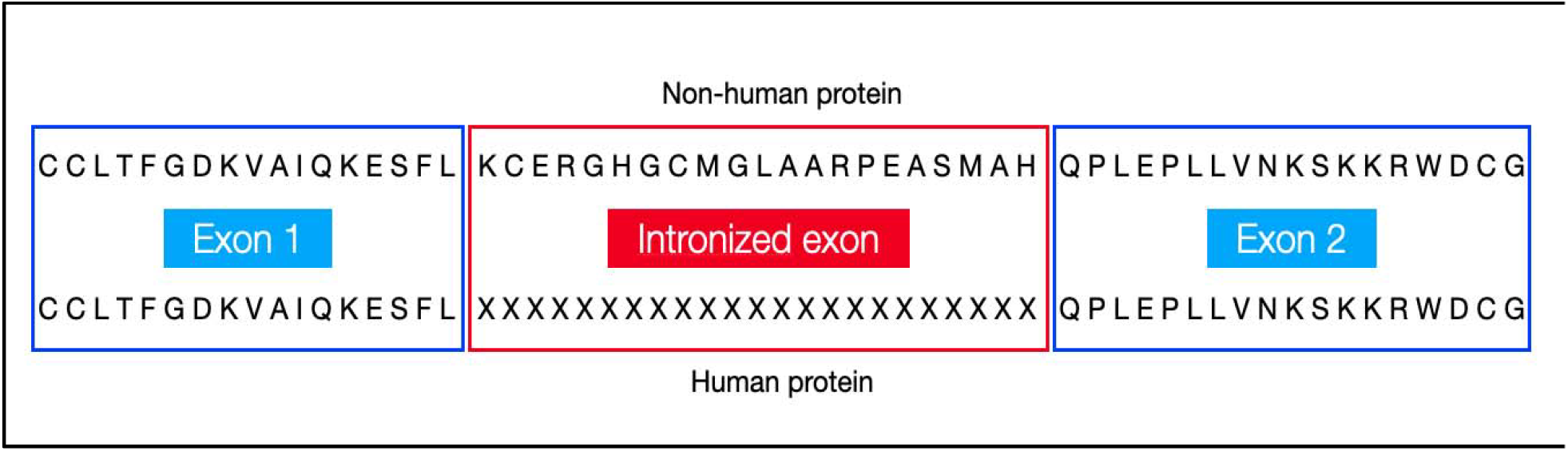
Alignment of sequences in a potential intronization event. We modified the human protein sequence (bottom) by inserting Xs at the position of the intron, as shown, and the modified sequence was aligned to all possible orthologs. The upper part of the figure shows an aligned sequence from an orthologous protein in another species. The ortholog contains additional amino acids spanning the position that is now an intron in humans.

To further investigate intronization events, we extracted the DNA sequence of the human intron, translated it in all three frames, and assessed its similarity to the orthologous proteins. Even though the human sequence is no longer constrained to encode a protein, some amino acid similarity might still be detectable. Additionally, we used ColabFold (Mirdita et al. 2022; G. Kim et al. 2023) to predict the structures of both the human and orthologous sequences and to compute confidence scores (pLDDT) for both. The predicted structures were then visualized in PyMOL.

## RESULTS

From the set of 19,120 MANE genes, we identified 584 recently gained introns in 514 distinct proteins. For each human gene that appeared to have gained an intron, we arranged the gene and its orthologs according to their taxonomic relationships as shown in **Figure 3**, where each intron is displayed with a small red square if present and a blue square if absent. Intron gain events can be identified by finding a vertical blue line connecting multiple genes, all of which are missing the intron, that surrounds a region where the intron is present, shown in red. As the figure shows, when we find a set of adjacent species in the tree that contain an intron, and those species are surrounded by others that are missing the intron, this finding corresponds to a subtree (containing human) that has gained an intron. We estimated the approximate origin of intron gain by identifying the subtree that maximizes the number R of aligned (red) introns and minimizes the number (B) non-aligned (blue) ones; i.e., that maximizes the quantity R-B.

**Figure 3.**
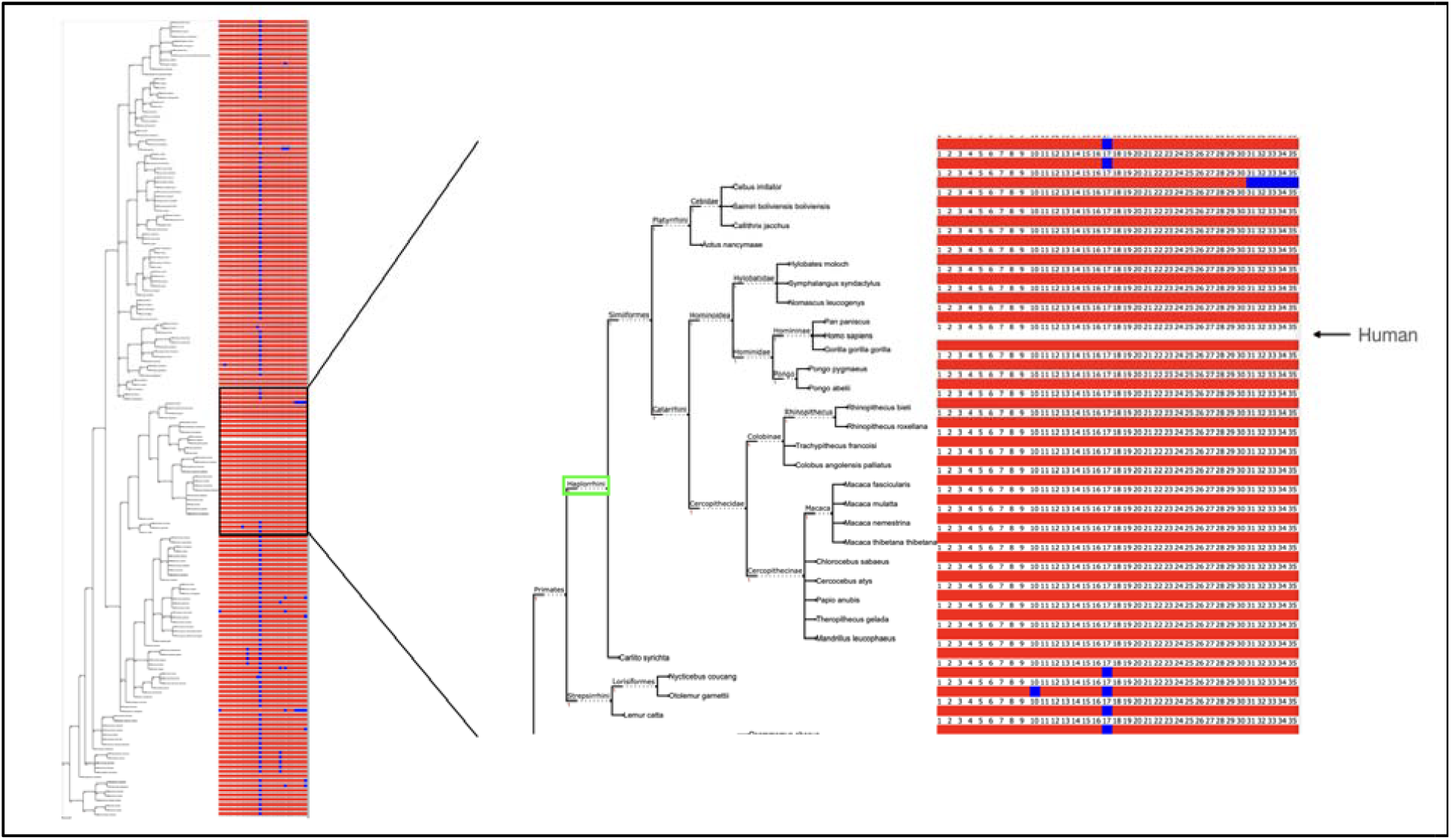
Phylogenetic analysis showing intron gain in gene A2M (GenBank accession NP_000005.3). The left panel shows a taxonomy of species that contain orthologues of A2M, with each species’ introns shown as a row of red or blue squares corresponding to the human gene, which has 35 introns. The squares to the right of each gene are colored red for introns that occur in the same position as the human sequence, and blue for introns present in humans but missing from the ortholog. The right panel provides a magnified view, highlighting the suborder Haplorhini in which the gain of intron 17 occurred.

**Figure 3** illustrates the gain of intron 17 in the gene A2M (Alpha-2-Macroglobulin) in humans and other members of the suborder Haplorhini, a suborder contained within the primates. Supplementary **Figures S2-S4** show three additional intron gain events, and a complete list of the 584 intron gains, including gene accession numberss, genomic coordinates, and taxonomic groups in which the intron gain event occurred, is provided in Supplementary Table S1.

From the 584 intron gain events, we identified three examples of intronization among all recently gained intron events as described in Methods. Although these represent a very small fraction of all events, they nonetheless appear to provide support for the hypothesis that intronization accounts for at least some intron gains. The three events we identified occurred in the genes CYP21A2, RPGR, and HELZ2. Each of these events features well-aligned flanking exons, highlighting conserved regions in the proteins. In addition, orthologous proteins in other species contain insertions of amino acid sequences at positions corresponding precisely to location of the human intron, and the lengths of those insertions match the length of a translation of the human intron, as shown by sequence alignments in **Figure 4** and Supplementary **Figures S4-S5**.

**Figure 4.**
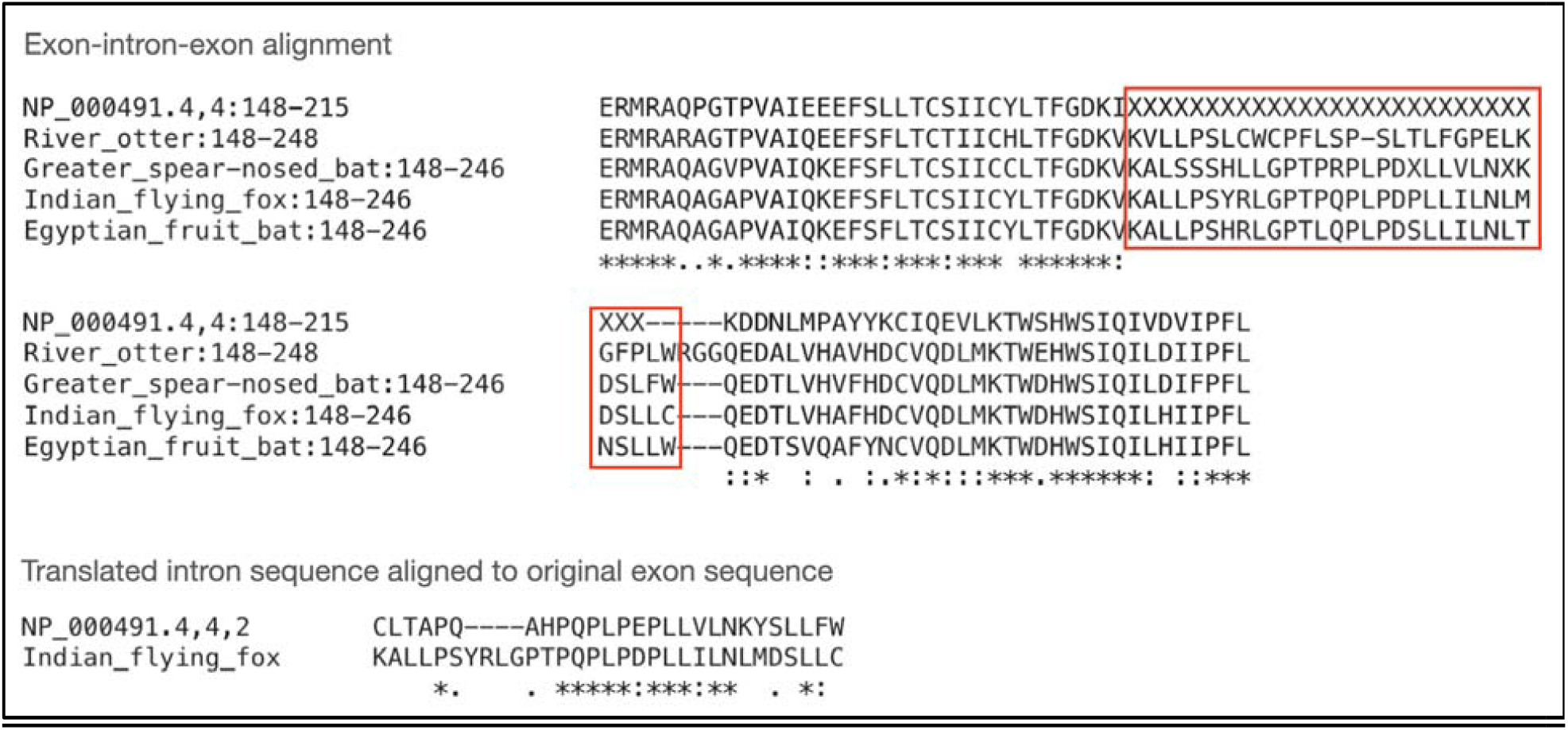
Evidence of intronization in intron 4 of CYP21A2 (NCBI accession NP_000491.4). Shown here is an alignment between the human protein and its orthologs in river otter (XP_032703087.1), greater spear-nosed bat (XP_045682190.1), Indian flying fox (XP_039738891.1), and Egyptian fruit bat (XP_036094280.1). The flanking exons on either side are very highly conserved. The X’s in the human protein were added in the position of intron 4, with the number of Xs corresponding to 1/3 of the length of the intron in nucleotides. As shown here, the orthologous proteins contain amino acid sequences that are similar in length to the expected sequence based on the human intron. The lower alignment demonstrates the similarity between the translated intron sequence in human (frame 2) and the original exon sequence in the Indian flying fox ortholog, which has a BLAST E-value of 5x10^-10^.

For each of these three proteins that had undergone intronization, we selected the orthologous proteins from the closest relative that did not contain the specific intron, and compared the translated intron sequence to the translations of the corresponding exon sequences in the orthologs. For CYP21A2 (NP_000491.4), we found similarity between the translated intron and orthologs, as shown in **Figure 4**. The other two cases showed no detectable sequence similarities between the human intron and the corresponding amino acid sequences in other species, possibly because the human intronic sequence had simply accumulated too many mutations.

We used ColabFold (Mirdita et al. 2022; G. Kim et al. 2023) to predict the structures of the three human proteins that had undergone intronization (CPY21A2, RPGR, and HELZ2) and their orthologous counterparts that retain the original exons. The protein structures for human CPY21A2 and one of its orthologs are shown in **Figure 5**, and those for RPGR and HELZ2 are shown in Supplementary **Figures S7-S8**. Interestingly, we observed an improvement in average pLDDT scores for each of the human protein sequences. Upon closer examination, we found that the intronized segments in the non-human proteins are predicted to form unstructured coil regions. This observation suggests that the intronization of these exons did not compromise the protein’s functional integrity.

**Figure 5.**
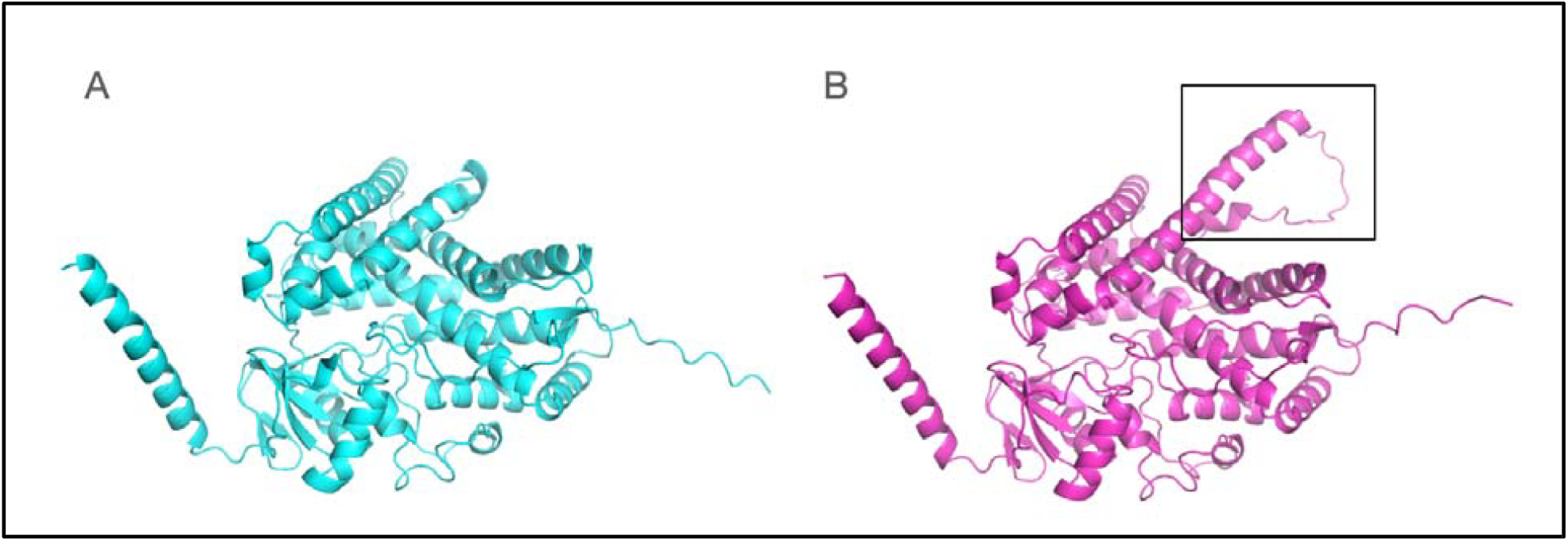
Protein structures of human protein CYP21A2 (NP_000491.4) (left) and its homolog in flying fox (XP_039738891.1) (right), as predicted by ColabFold. Notably, the region in the flying fox protein that aligns with the intronized sequence in human, indicated by a box, is mostly an unstructured coil region, which lowers the overall pLDDT score and suggests that it is not needed for the protein to function. The human protein’s pLDDT score is 90.2, while the fox protein’s pLDDT score is 85.9.

### In-depth analysis of potential recently-gained introns

We mapped intron gain events to divergence times and discovered that a majority of these events occurred more recently in evolutionary history, with a significant portion happening after the emergence of primates (Figure 6). This observation might be a consequence of our methodology, which relies on alignment and is therefore inherently more reliable at detecting recent events. We looked at the lengths of recently gained introns and found they were similar to all intron lengths in the MANE dataset (Figure 7), suggesting that the length of an intron does not influence the likelihood of its gain. Interestingly, our method detected a slight tendency for recently gained introns to appear near the ends of proteins (Figure 7C). This could be indicative of these positions being less stable, although further research is required to confirm this hypothesis. Additionally, we used PANTHER (Mi and Thomas 2009) to categorize proteins with recently gained introns based on their functions, but this analysis did not reveal any particularly notable trends.

**Figure 6.**
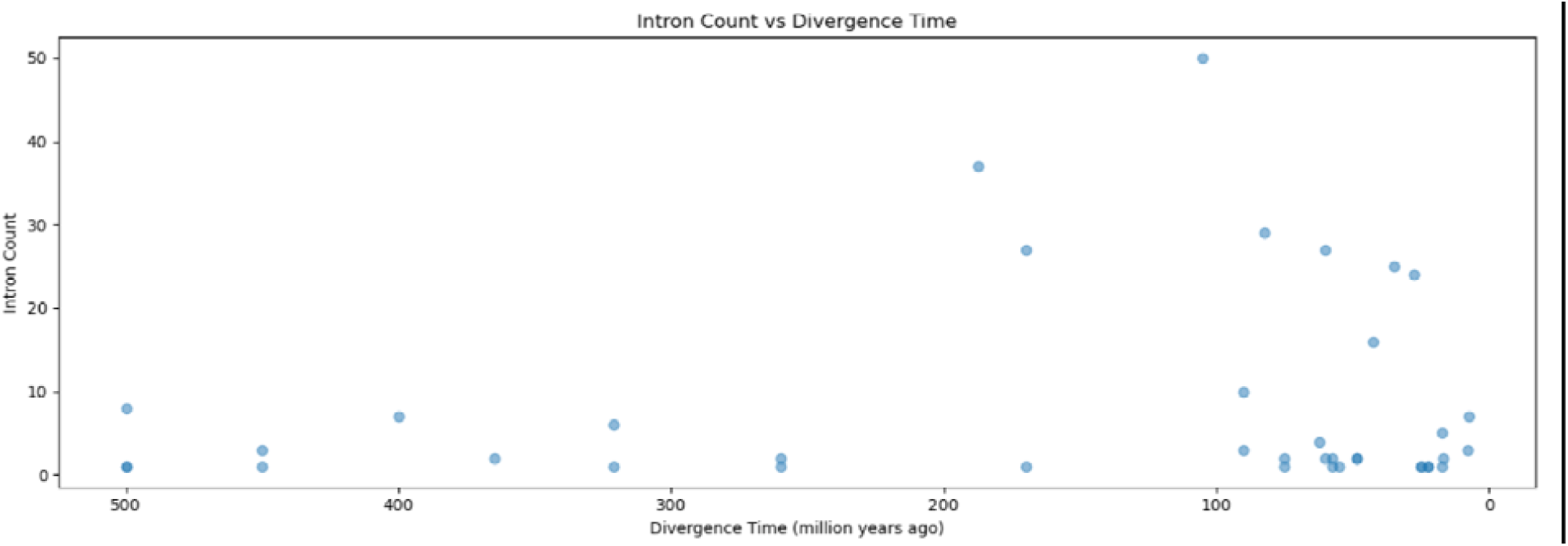
Number of introns gained in humans as a function of divergence time for intron gain events found in this study. The y-axis represents the number of events detected and the x-axis indicates the estimated time since divergence.

**Figure 7.**
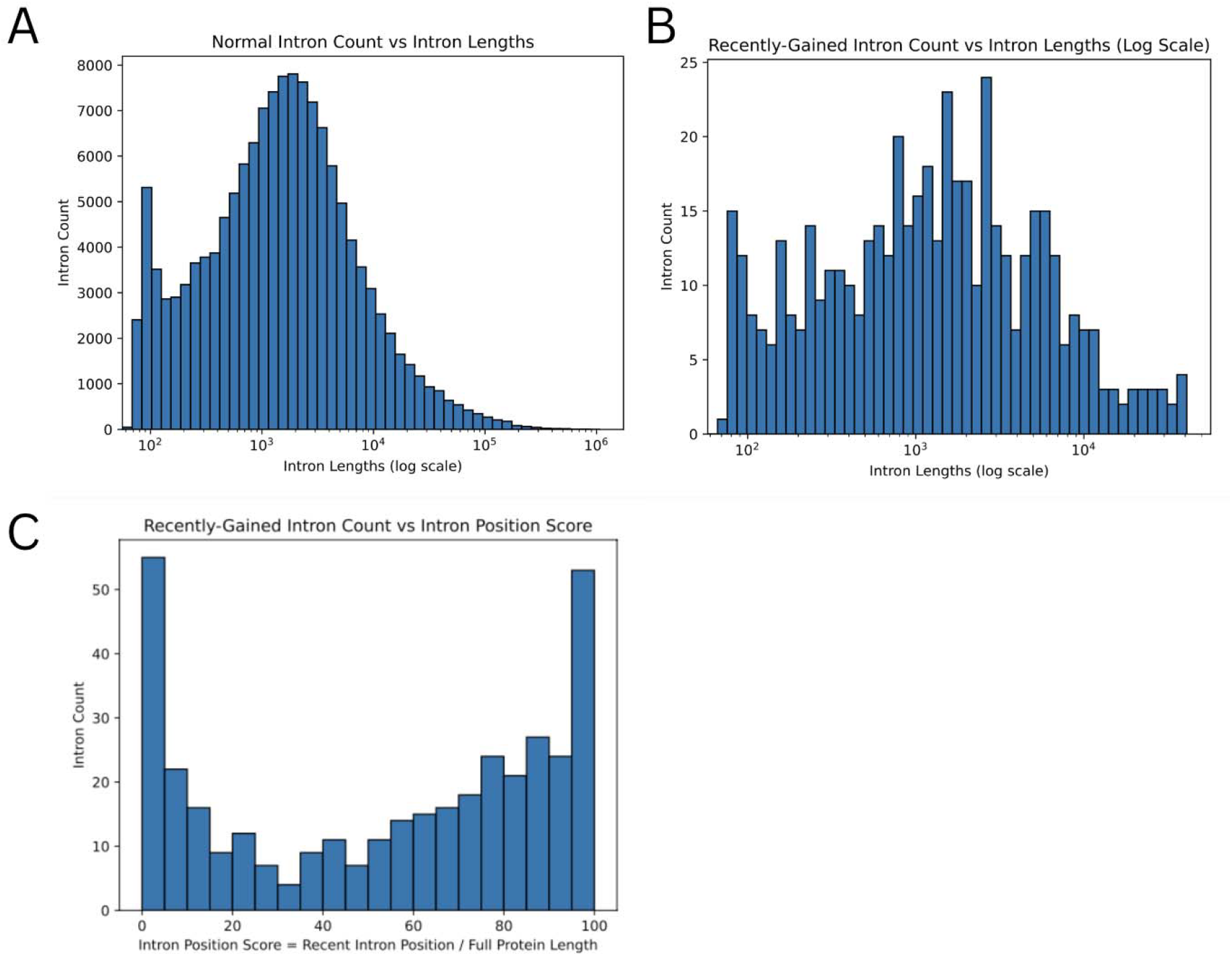
Histograms of intron lengths and positions for intron gain events. (A) The distribution of intron lengths of all MANE human introns. (B) Length distribution of recently gained introns. (C) Counts of recently gained introns plotted against their relative position in the protein.

## DISCUSSION

Earlier studies suggested that intron gains within a specific lineage are rare (Scott W. Roy, Fedorov, and Gilbert 2003), but become more frequent when comparing across different evolutionary subgroups (Carmel et al. 2007). Fedorov and colleagues reported in 2003 that only about 14% of animal introns align with plant intron positions, although that study was based on far less data than is available today (Fedorov et al. 2003). Our study used more than 17 million proteins spanning over 3400 vertebrate species to successfully identify numerous instances of intron gains in the vertebrate lineage, focusing specifically on gains in human genes. In addition to casting new light on the origins of introns, our findings may also benefit genome annotation. The most common strategy for annotating genes today is to align the known genes from other species, using both DNA-based and protein-based alignments. If an intron is not shared among species, then it might be missed when using this strategy.

As a mechanism for intron gain, intronization appears to be relatively rare, at least according to the methods described here. While indirect evidence for intronization events has been identified within several species, including *Cryptococcus, Caenorhabditis, Plasmodium vivax*, as well as in primate and rodent retrogenes (Irimia et al. 2008) (Scott W. Roy 2009) (Szcześniak et al. 2011) (Yang and Huang 2011), we have not encountered any studies that report intronization events across different subgroups of vertebrates. Identifying intronization events can be challenging due to sequence divergence and potential changes in length resulting from mutations with introns. However, the three cases we identified are particularly compelling. In these instances, the lengths of the exon sequences in other species closely match that of the translated intronized sequences in humans. Moreover, the intronized portions appear to preserve or even enhance their respective protein structures, as measured by the pLDDT scores assigned by AlphaFold2.

## ACKNOWLEDGEMENTS

This work was supported in part by NIH grants R01-HG006677 and R35-GM130151.

