## Supplementary File for "Discovering Intron Gain Events in Humans through Large-Scale Evolutionary Comparisons"

**Supplementary materials**

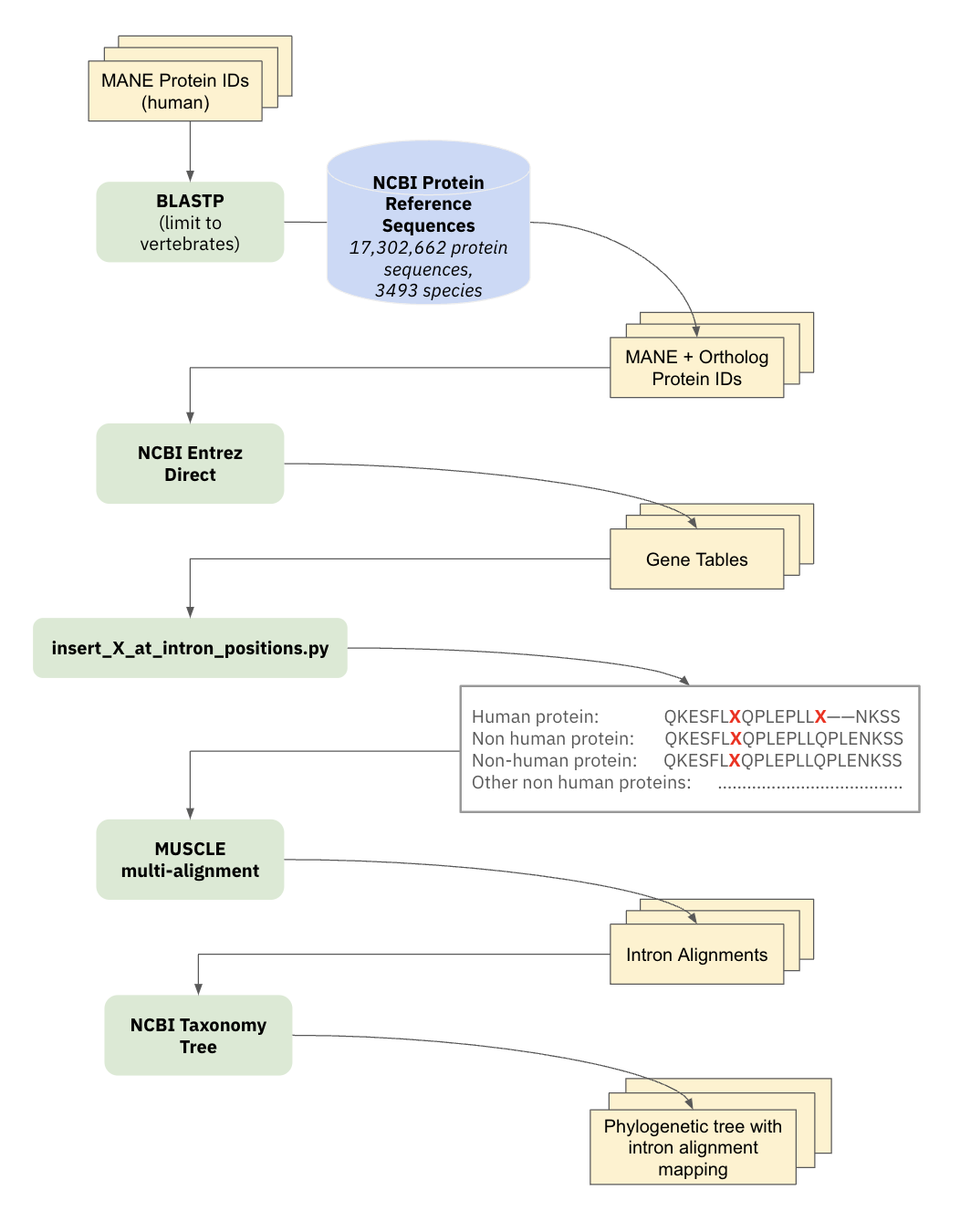

**Supplementary Figure S1**. Overview of the pipeline for detecting intron gain events in human proteins. The workflow begins with the input of a protein sequence from MANE, followed by a BLASTP search to obtain orthologous proteins from other vertebrates. Next, the intron positions and lengths are computed using gene tables retrieved from NCBI via Entrez Direct. Each intron position is marked with an ‘X’ inserted into the amino acid sequence of all proteins containing that intron. The sequences and the ‘X” placeholders are then aligned using MUSCLE. For each intron in each human protein, the pipeline looks for corresponding introns in the orthologous proteins. Finally, the NCBI taxonomy is employed to assign scores to different taxonomic subgroups, allowing us to infer the evolutionary point of intron gain by identifying the subgroup that maximizes the number of aligned introns and minimizes the number of missing introns.

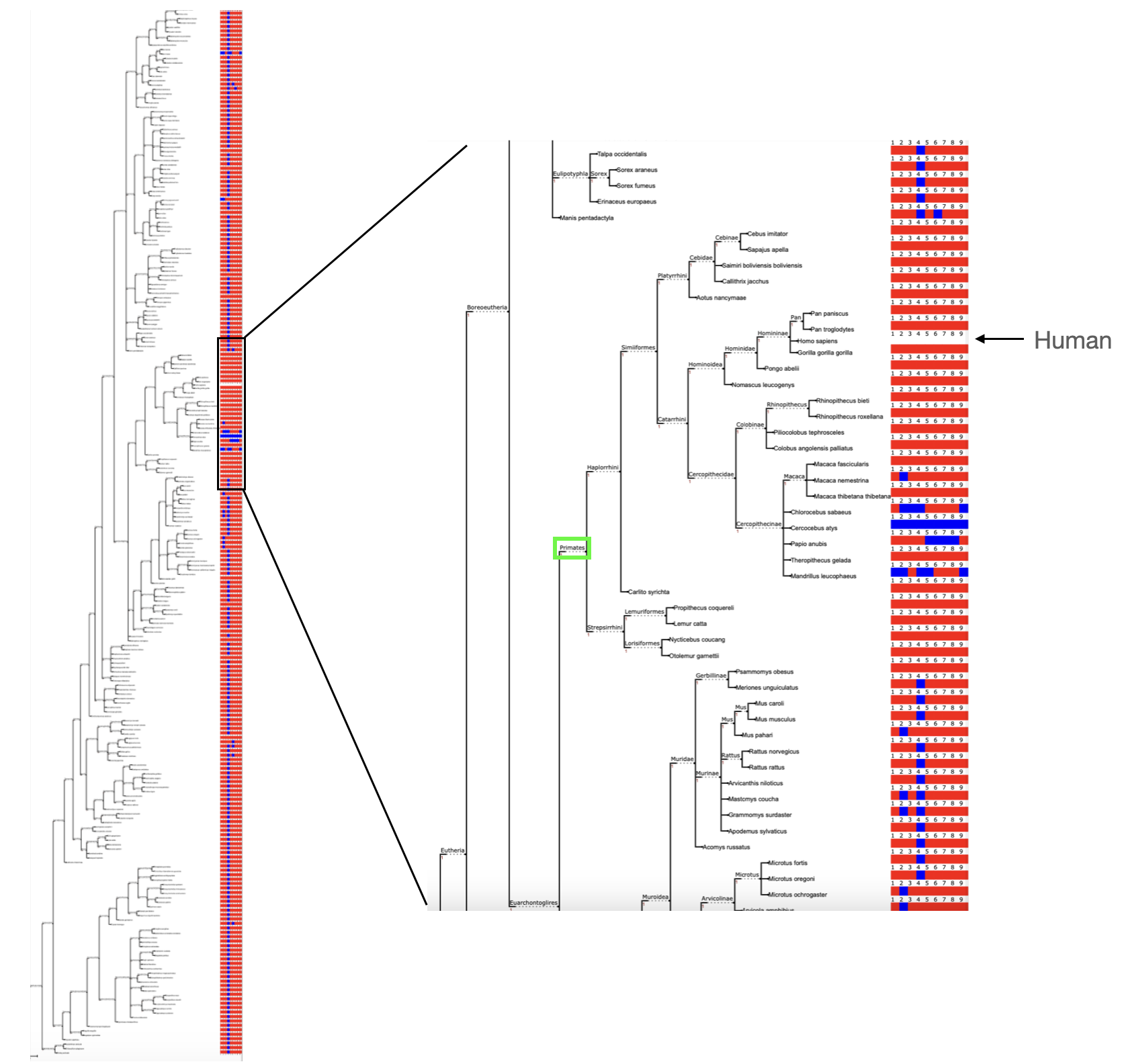

**Supplementary Figure S2**: Phylogenetic analysis showing intron gain in CYP21A2 (GenBank accession NP_000491.4). The left panel shows a taxonomy of species that contain orthologues of CYP21A2, with each species’ introns shown as a row of red or blue squares corresponding to the human gene, which has 9 introns. The squares to the right of each gene are colored red for introns that occur in the same position as the human sequence, and blue for introns present in humans but missing from the ortholog. The right panel provides a magnified view, highlighting the suborder Primates in which the gain of intron 4 occurred.

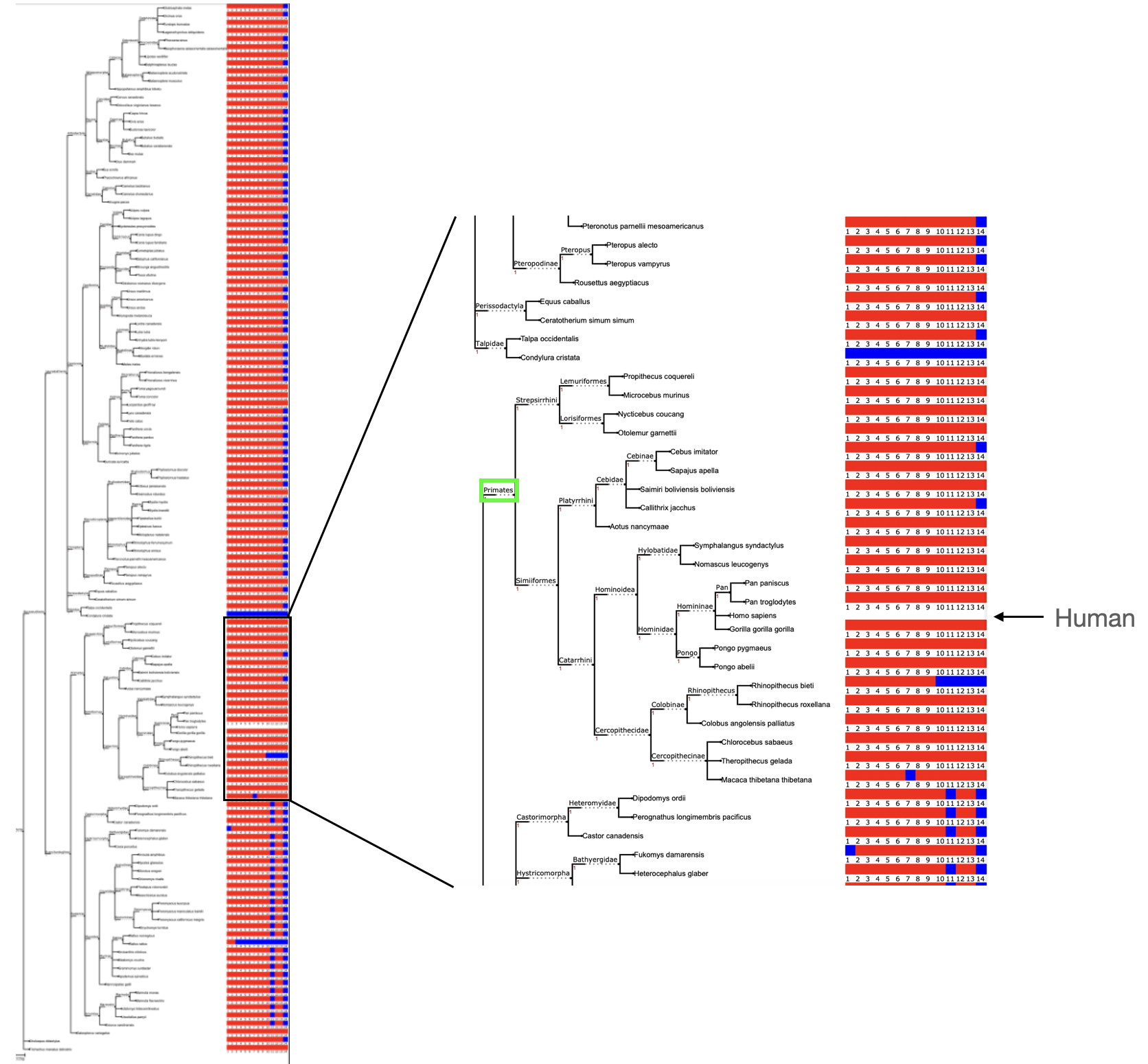

**Supplementary Figure S3**: Phylogenetic analysis showing intron gain in RPGR (GenBank accession NP_001030025.1). The left panel shows a taxonomy of species that contain orthologues of RPGR, with each species’ introns shown as a row of red or blue squares corresponding to the human gene, which has 14 introns. The squares to the right of each gene are colored red for introns that occur in the same position as the human sequence, and blue for introns present in humans but missing from the ortholog. The right panel provides a magnified view, highlighting the suborder Primates in which the gain of intron 14 occurred.

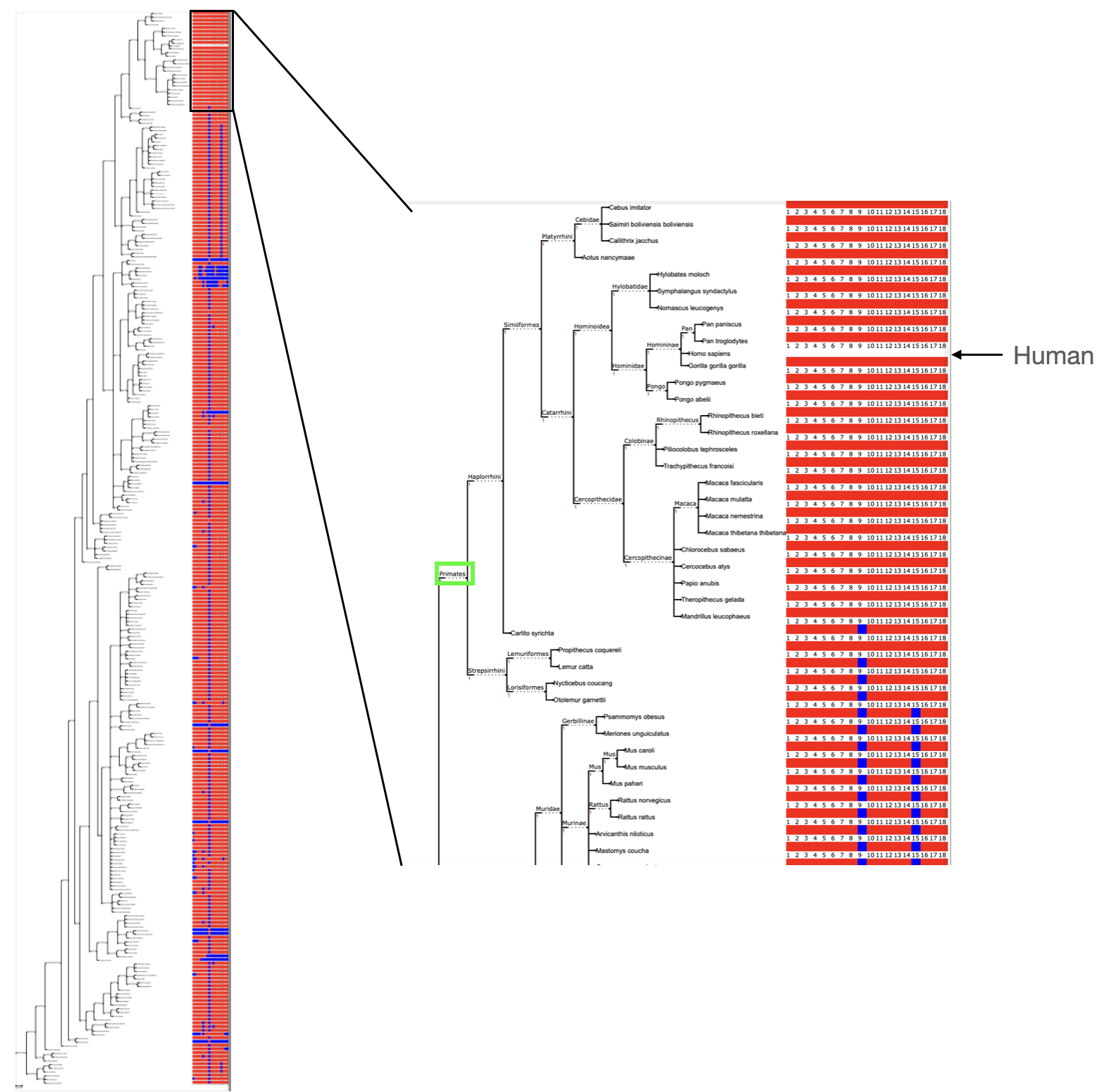

**Figure S4**: Phylogenetic analysis showing intron gain in HELZ2 (GenBank accession NP_001030025.1). The left panel shows a taxonomy of species that contain orthologues of HELZ2, with each species’ introns shown as a row of red or blue squares corresponding to the human gene, which has 18 introns. The squares to the right of each gene are colored red for introns that occur in the same position as the human sequence, and blue for introns present in humans but missing from the ortholog. The right panel provides a magnified view, highlighting the suborder Primates in which the gain of intron 9 occurred.

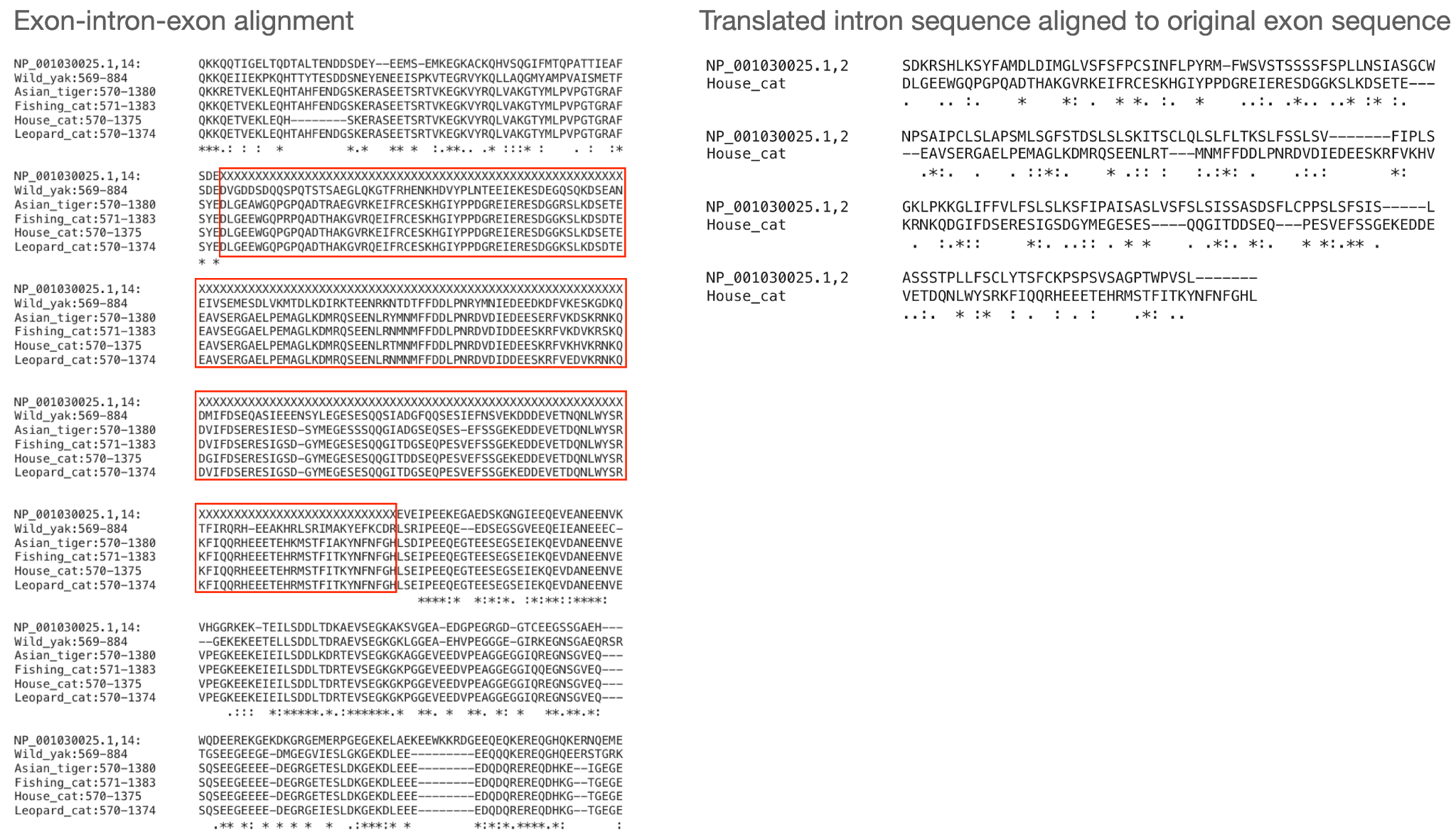

**Figure S5**. Evidence of intronization for intron 14 of human protein RPGR (NCBI accession NP_001030025.1). Shown here is an alignment between the human protein and its orthologs in wild yak (XP_005889507.2), Asian tiger (XP_042830144.1), fishing cat (XP_047699385.1), house cat (XP_044907220.1), and (leopard cat XP_043426382.1). The human intron, indicated in red boxes, is represented by X's, and the orthologous proteins contain amino acid sequences that are almost exactly as long as the expected sequence based on the human intron.

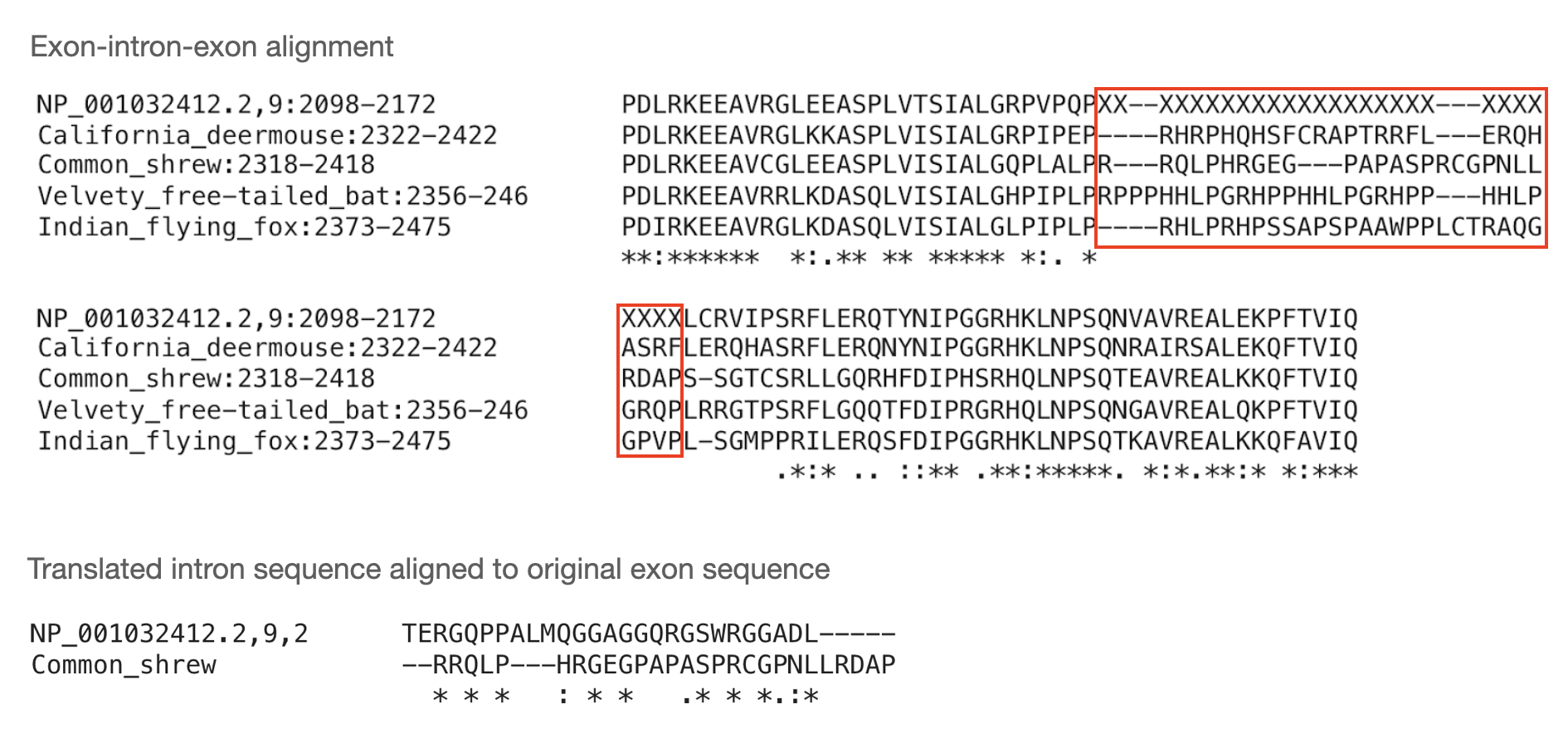

**Figure S6.** Evidence of intronization in HELZ2 (NCBI accession NP_001032412.2), intron 9. Shown here is an alignment between the human protein and its orthologs California deer mouse XP_052579411.1, common shrew XP_054994474.1, velvety free-tailed bat XP_036099976.1, and Indian flying fox XP_039720752.1. The orthologous proteins contain an amino acid sequence that is almost as long as the expected sequence based on the human intron.

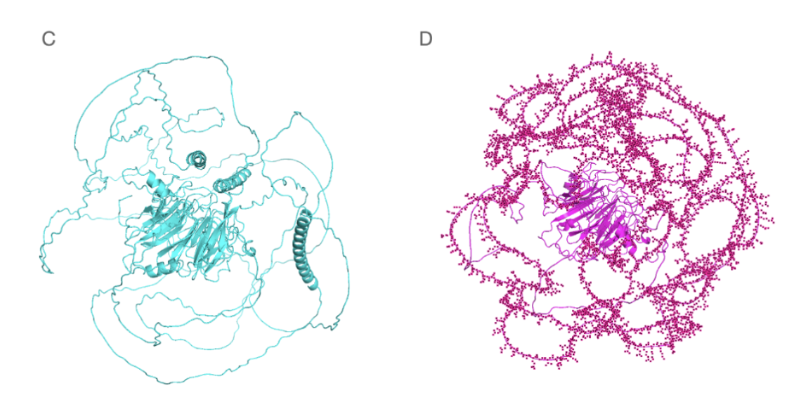

**Supplementary Figure S7**: Protein structures of human RPGR protein NP_001030025.1 (green) and house cat protein XP_044907220.1 (blue), as predicted by AlphaFold2 and ColabFold. Notably, the region in XP_044907220.1 that aligns with the intronized sequence in NP_001030025.1 is depicted as an unstructured coil region, which lowers the overall score assigned by AlphaFold2 and suggests that it is not needed for the protein to function. Both proteins get low scores overall: the human protein's pLDDT score is 52.2 and the cat protein's pLDDT score is 47.5.
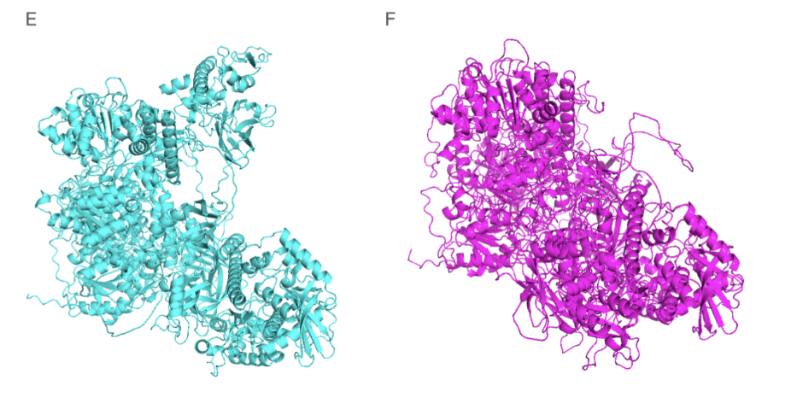
 **Supplementary Figure S8**: Protein structures of human HELZ2 protein NP_001032412.2 (green) and common shrew protein XP_054994474.1 (blue), as predicted by AlphaFold2 and ColabFold. The region in XP_054994474.1 that aligns with the intronized sequence in NP_001032412.2 is an unstructured coil region, which lowers the overall score assigned by AlphaFold2 and suggests that the additional sequence in the shrew protein might not be not needed.
